## Supplemental Figures with Legend for "Proximity Labeling of NIMA Kinase Complex Components in *C. elegans*"

S1 Fig

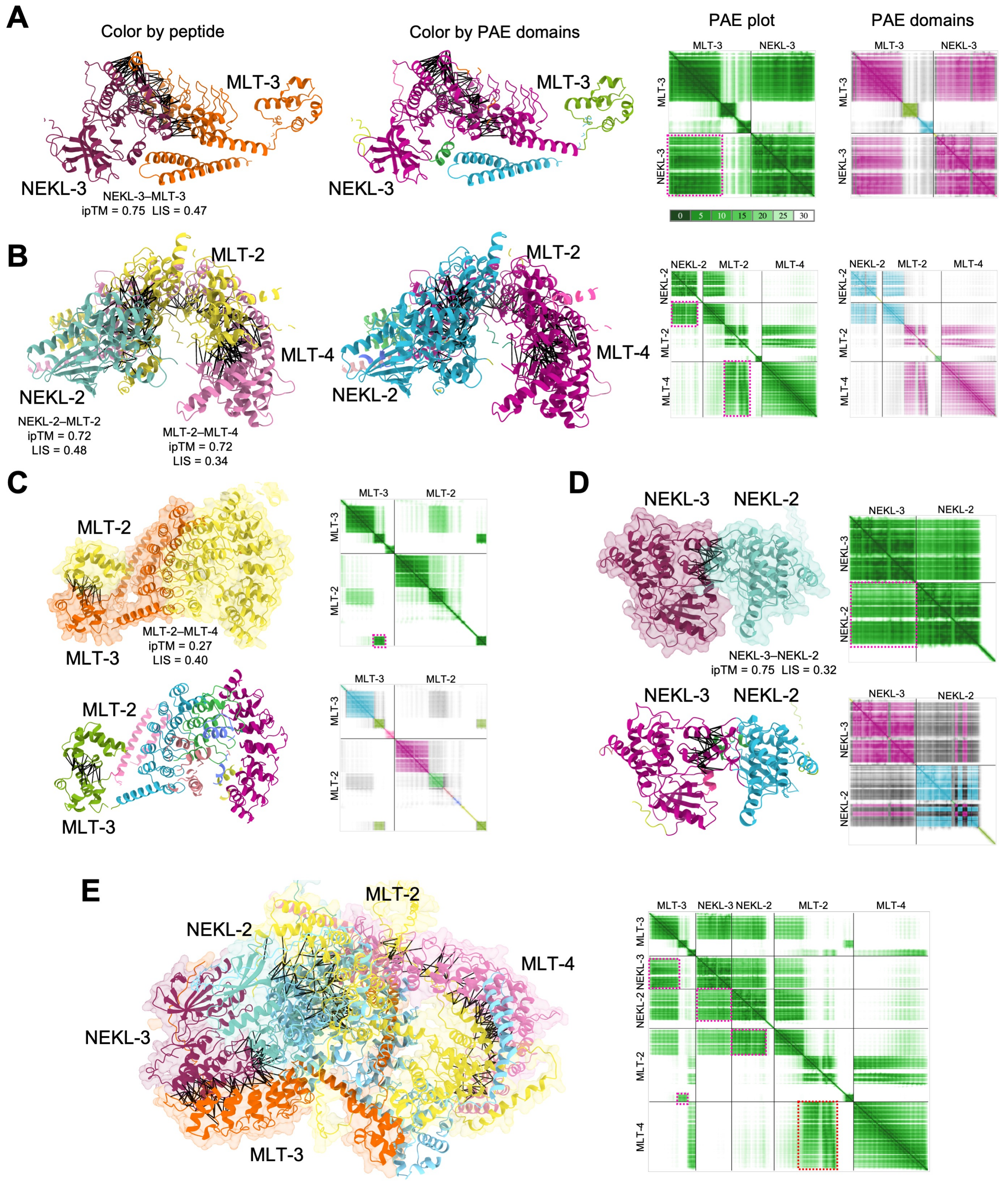


**S1 Fig. AlphaFold modeling of NEKL–MLT interactions.** (A–E) AlphaFold3 modeling and ChimeraX rendering of predicted (A) NEKL-3–MLT-3, (B) MLT-2–MLT-4–NEKL-2, (C) MLT-2–MLT-3, (D) NEKL-2–NEKL-3, and (E) MLT-3–NEKL-3–NEKL-2–MLT-2–MLT-4 protein complexes. Black lines between proteins indicate high-confidence Predicted Aligned Error (PAE) pseudobonds (PAE ≤ 6Å; interatomic-distance ≤ 5Å). For clarity, highly unstructured portions of the peptides

(B-factor ≤ 30) were hidden. Panels A–E show protein-backbone ribbon diagrams (colored by peptide) along with single-color (green) PAE diagrams indicating regions of high structural confidence (low PAE) within and between the indicated proteins. Dashed-line cyan boxes correspond to regions where the two proteins are predicted for form a coherent structural unit. Panels A–D additionally show protein ribbon diagrams colored by PAE domain along with corresponding color-matched PAE diagrams. Ribbon diagrams in panels C–E also show partially transparent surface renderings**.** Inter-chain predicted Template Modeling (ipTM) values, and Local Interaction Scores (LIS) are shown for the indicated proteins modeled as heterodimers by AlphaFold3. Note that the five-protein complex (panel E) predicts very similar protein–protein interfaces as the subcomplex models, consistent with the hypothesis that the two subcomplexes can physically interact.

### S2 Fig

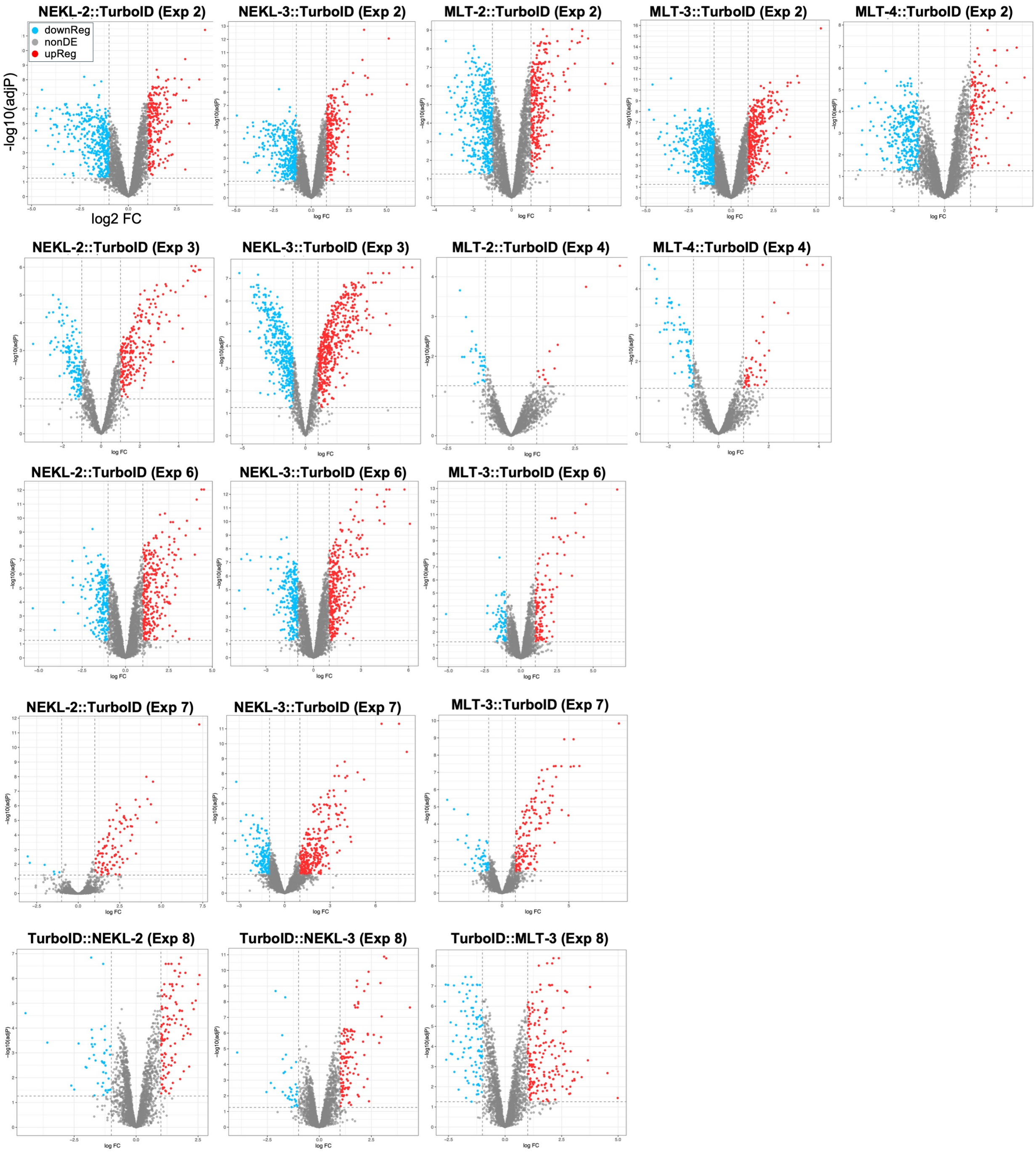


**S2 Fig. Volcano plots.** Volcano plots of bait–TurboID strains versus N2 control showing the $\log_2$ fold change (FC) versus $-\log_{10}$ adjusted p-value. Red and blue dots indicate proteins that were upregulated and downregulated, respectively, in the bait strain relative to the N2 strain; select identified bait and prey proteins are indicated by arrows. NonDE, not differentially expressed. Red and blue dots indicate proteins that were upregulated in the bait and N2 strains, respectively. Additional volcano plots are shown in Fig 3A, 6B, S4 Fig, and S5 Fig. Data corresponding to volcano plots are available in S1–8 Files.

### S3 Fig

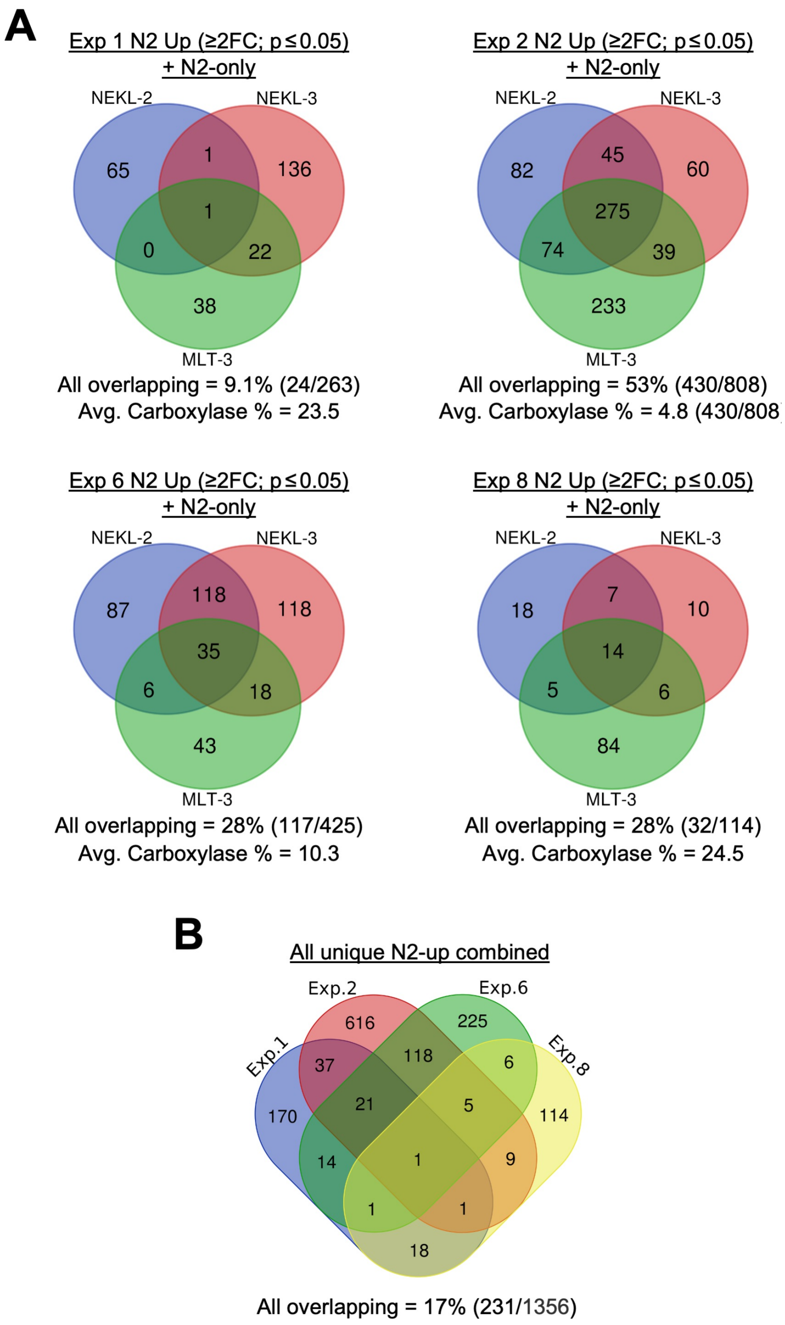


**S3 Fig. Analysis of N2 upregulated proteins.** (A) Venn diagrams showing overlap of proteins upregulated (≥2-fold (FC); adjusted p ≤ 0.05) in N2 samples (N2 Up) versus NEKL-2::TurboID, NEKL-3::TurboID, and MLT-3::TurboID bait samples from Exp 1, Exp 2, Exp 6, and Exp 8. N2-only indicates proteins that were present in all technical replicates of the N2 sample but were absent in all technical replicates of the corresponding bait–TurboID; Also see Fig 2B. Note variability in the extent of overlap among experiments. (B) Venn diagram comparing overlaps among the combined N2 upregulated proteins from Exp 1, Exp 2, Exp 6, and Exp 8. Note the relative lack of consistently overlapping proteins among experiments.

S4 Fig

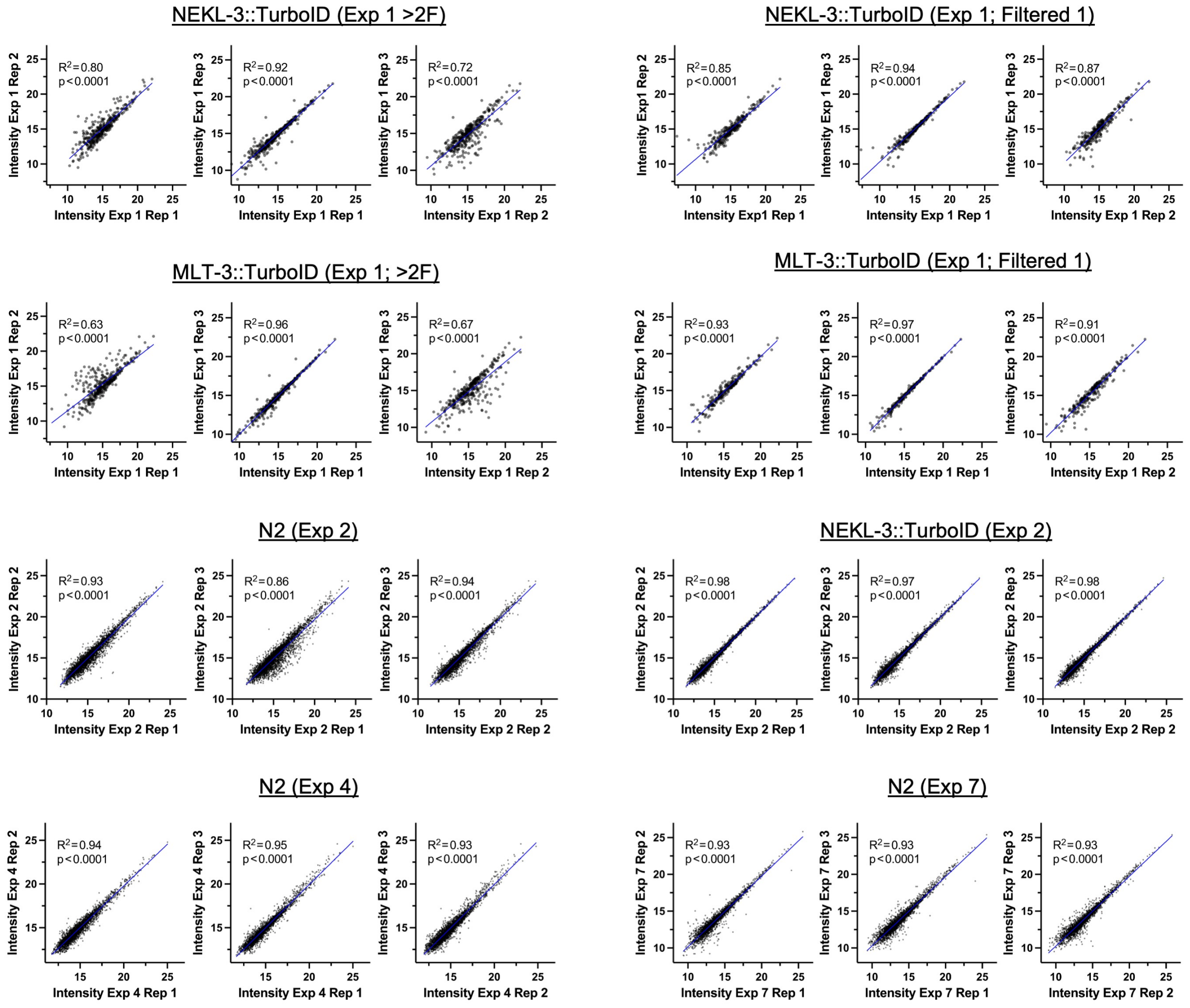


**S4 Fig. Additional analysis of technical replicates.** Scatter plot diagrams showing correlations of individual protein intensities (abundance) between the indicated replicate pairs. $R^2$ values were derived using simple linear regression; raw data are available in S10 File. Also see Fig 5.

S5 Fig.

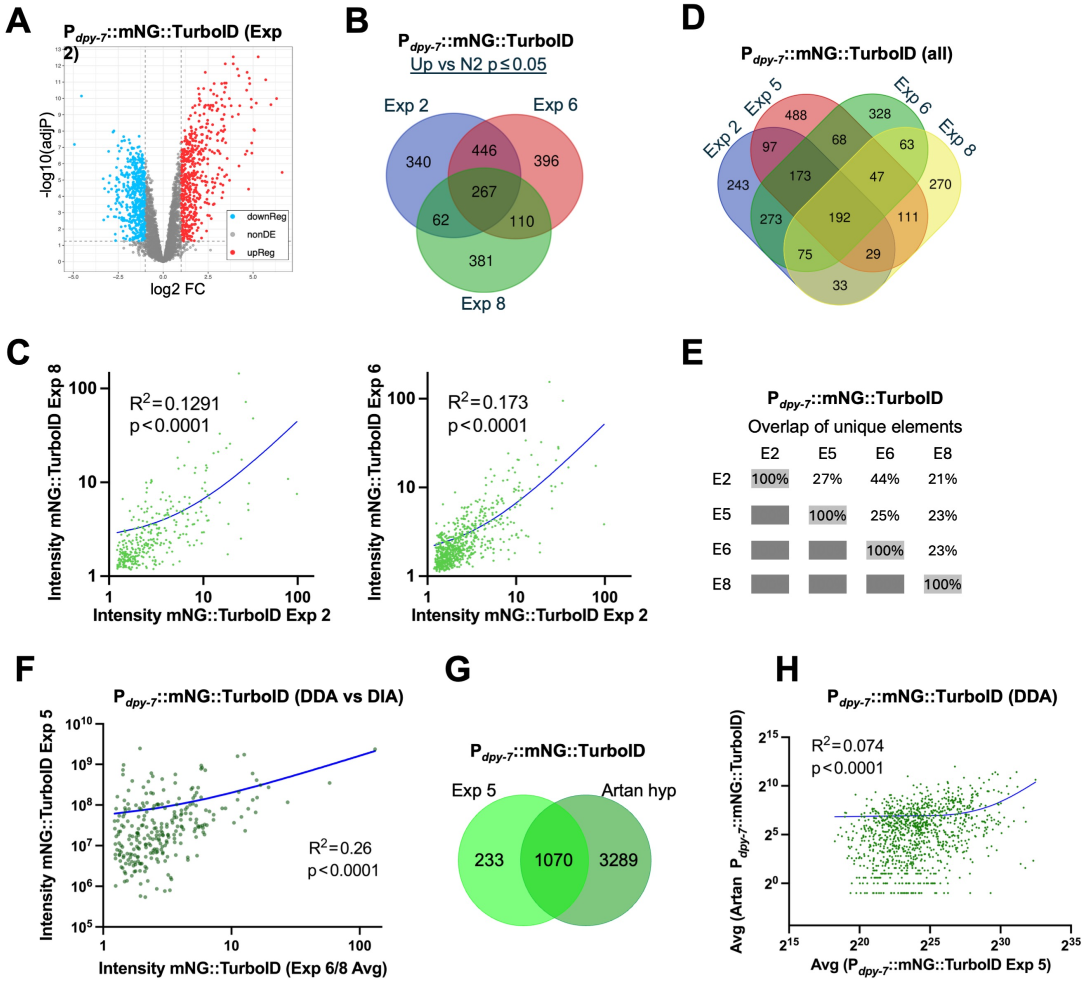


**S5 Fig. Additional analysis of P$_{dpy\text{-}7}$::mNG::TurboID datasets.** (A) Volcano plot of P$_{dpy\text{-}7}$::mNG::TurboID versus N2 showing $log_2$ fold change (FC) versus –$log_{10}$ adjusted p-value from Exp 2. (B) Venn diagram overlap of proteins significantly upregulated (based on adjusted p-values) in the three DIA-MS P$_{dpy\text{-}7}$::mNG::TurboID experiments. E, Experiment. (C) Scatter plots showing correlations of protein intensities (abundance) for the mNG::TurboID control strain from indicated experimental pairs. $R^2$ values were derived using simple linear regression; raw data are available in S11 File. The curved regression lines are due to the conversion of axes to a log scale. The markedly lower $R^2$ values, relative to Exp 6 vs Exp 8, are due in large part to discrepancies in intensity values at the high end of the scale. (D) Venn diagram and (E) quantification of pairwise overlaps for the four P$_{dpy\text{-}7}$::mNG::TurboID experiments. Proteins included were detected in all technical replicates. Note that although Exp 2 and Exp 6 showed the greatest overlap in detected proteins, the correlation in protein levels was strongest between Exp 6 and Exp 8 (also see Fig 6C). (F) Scatter plot showing weak correlation between Exp 5 (DDA-MS) and averaged data from Exp 6 and Exp 8 (DIA-MS). (G) Overlap and (H) correlation between Exp 5 and DDA-MS experiment from Artan and colleagues (PMID: 35665632). Note that although the proteins detected in Exp 5 overlapped extensively with a subset of those previously reported, the correlation in their expression levels was weak.

### S6 Fig

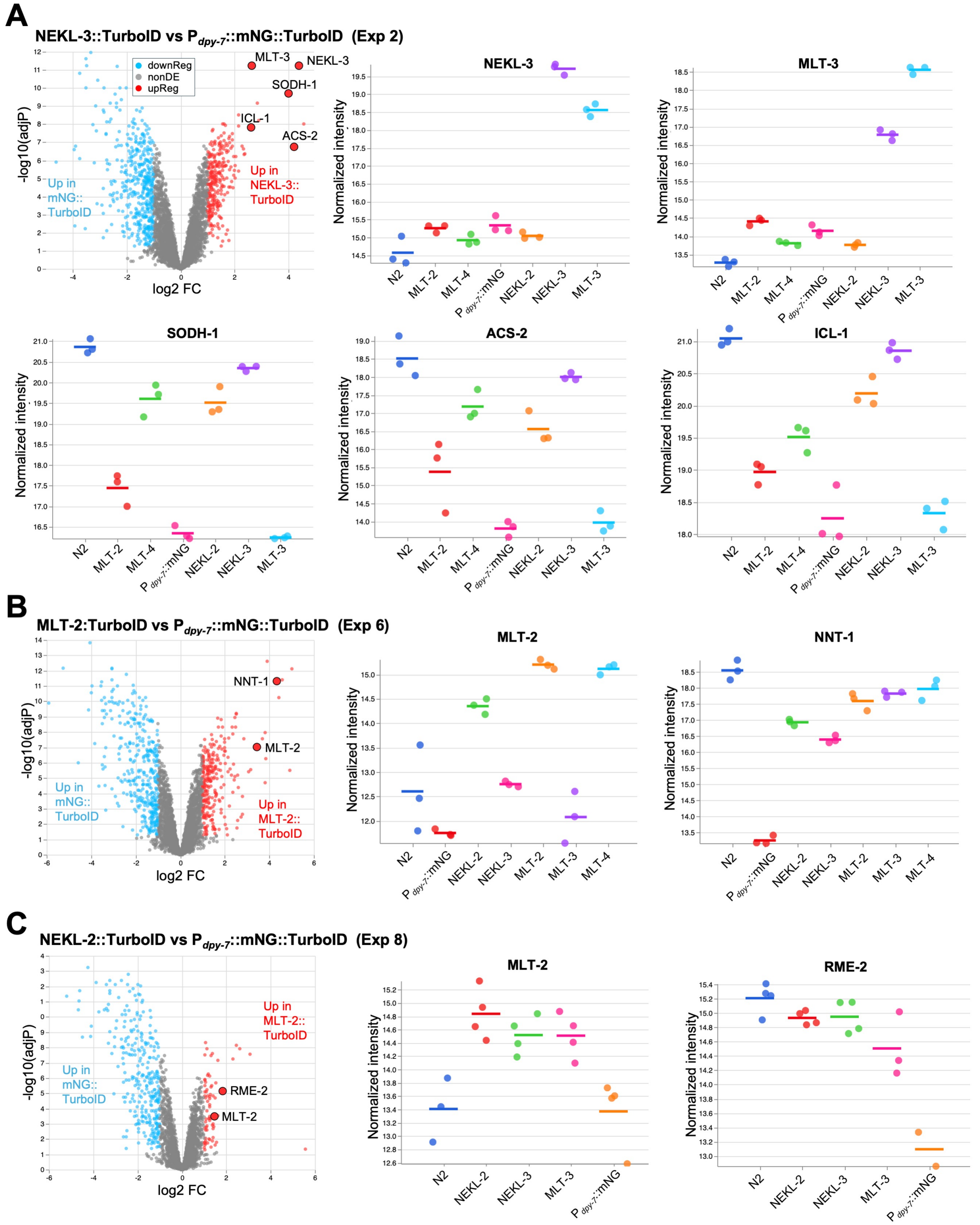


**S6 Fig. $P_{dpy\text{-}7}$::mNG::TurboID downregulated proteins.** (A–C) Volcano plots of $P_{dpy\text{-}7}$::mNG::TurboID versus NEKL-3::TurboID (A), MLT-2::TurboID (B), and NEKL-2 TurboID (C) indicating the $\log_2$ fold change (FC) versus $-\log_{10}$ adjusted p-values. Enlarged red dots indicate proteins shown in the accompanying intensity plots. Intensity plots ($\log_2$ scale) showing the abundance of the indicated bait and prey proteins; individual dots represent values from the three technical replicates. For SODH-1, ACS-2, ICL-1, NNT-1, and RME-2, levels in $P_{dpy\text{-}7}$::mNG::TurboID were markedly lower than in the N2 or most bait–TurboID strains.

### S7 Fig

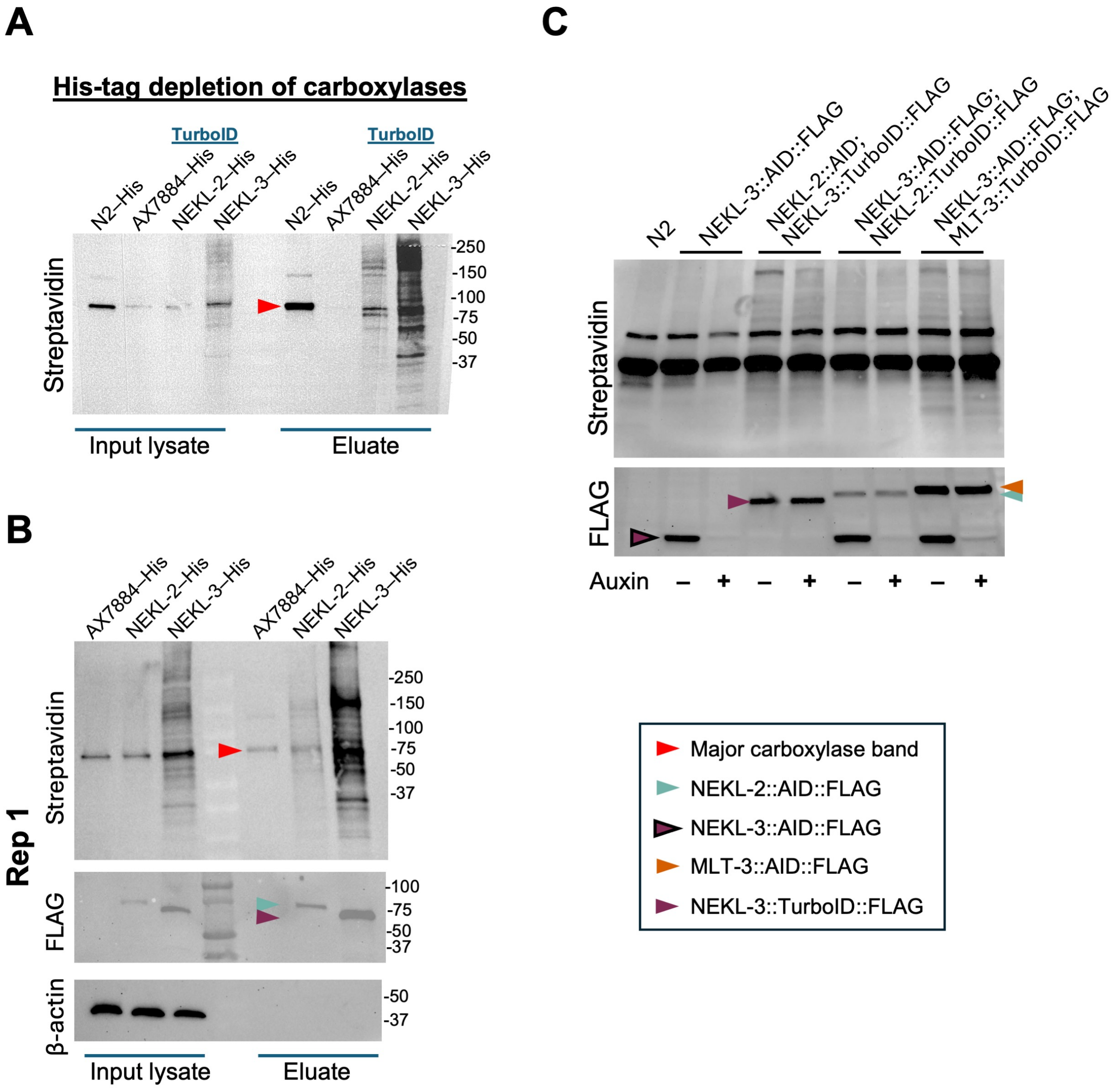


**S7 Fig. Methodological variations.** (A–C) Western blots with specific proteins indicated by the key. (A) Streptavidin-probed western blot showing a reduction in the major carboxylase band in strains that underwent carboxylase depletion (AX7884, NEKL-2, and NEKL-3). –His, lysates were incubated with nickel-coated (His-binding) beads to remove the endogenous His-tagged versions of PYC-1, PCCA-1, MCCC-1, and POD-2. N2 does not encode His-tagged carboxylases, accounting for the stronger band in this strain. Increased protein concentration in the eluate lanes facilitates the visualization of the bait-induced biotinylated bands. (B) Western blots from Exp 3, replicate 1Exp  showing biotinylated products (streptavidin), bait proteins (FLAG), and loading control (β-actin). (C) Western blot of lysates from the indicated strains after incubation in the presence or absence of auxin, which leads to the degradation of the AID-tagged proteins. Note that the NEKL-2::AID strain contained a defective FLAG tag and was thus not detectable.

S8 Fig

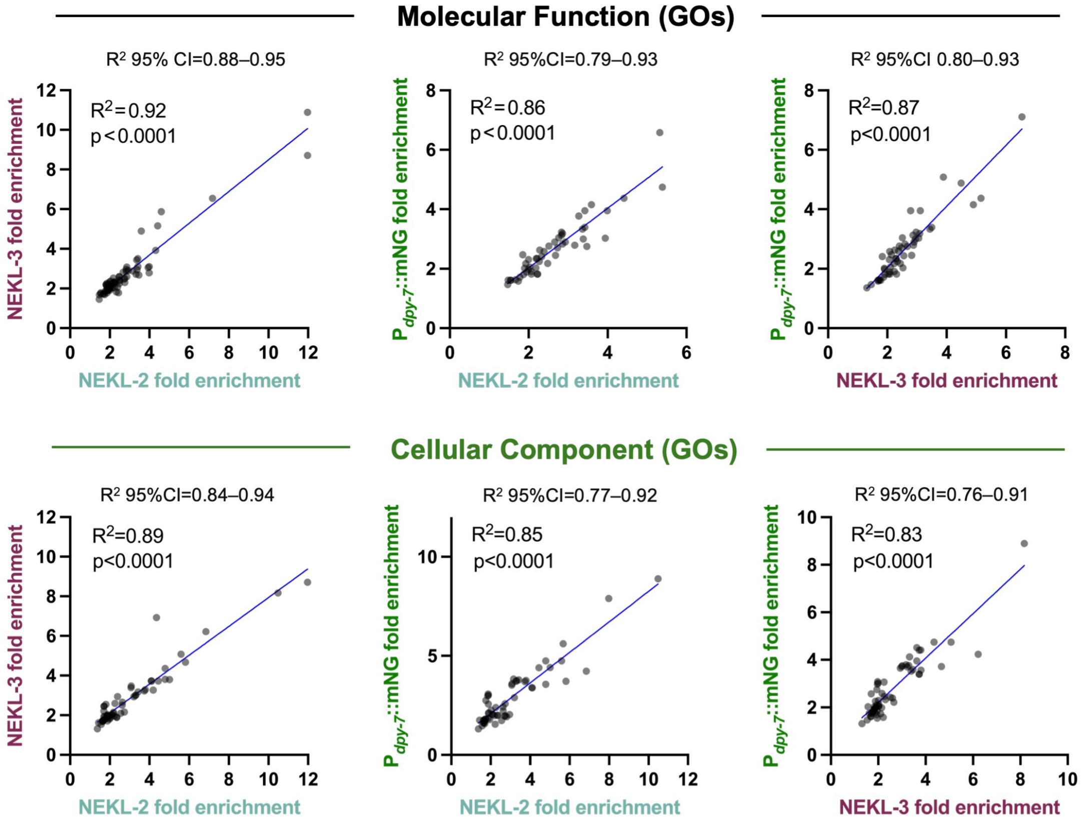


**S8 Fig. Correlations of GO terms**. Scatterplots showing correlations in fold enrichments of Molecular Function and Cellular Component GOs for the combined NEKL-2::TurboID, NEKL-3::TurboID, and $P_{dpy-7}$::mNG::TurboID datasets. 95%CI, 95% confidence interval. Although correlation was greatest between NEKL-2–NEKL-3, this was not statistically significant based on CIs.
