## Supplementary material for "Proximity Labeling of NIMA Kinase Complex Components in *C. elegans*": S1 Data

### S1 Data. Western blot data for Exp 1–8

#### Experiment 1

Rep 1

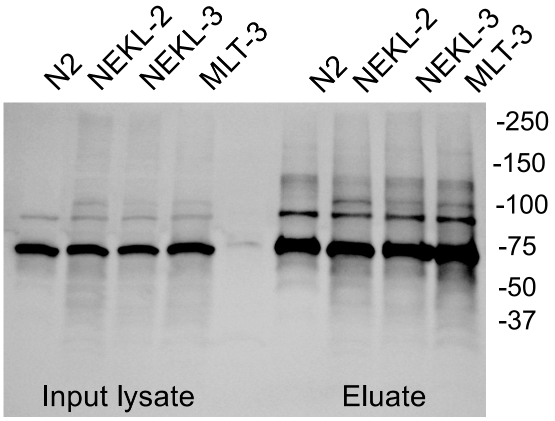


**Streptavidin**

Rep 2

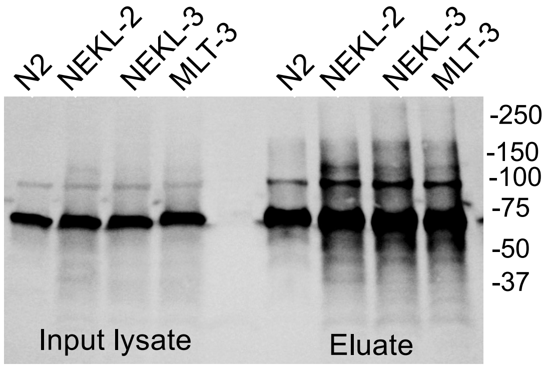


**Streptavidin**

Rep 3

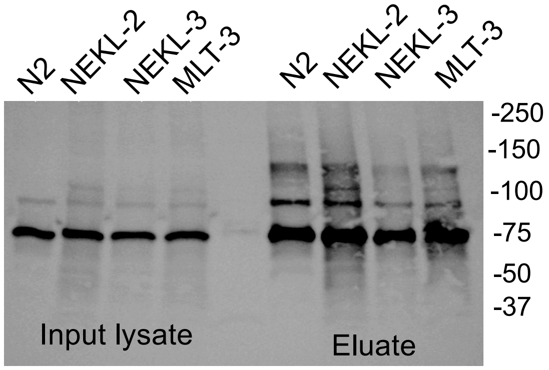


**Streptavidin**

PYC-1a, 129.3 kDa; PYC-1b, 67.8 kDa; PCCA-1, 79.7 kDa; MCCC-1, 73.7 kDa; POD-2a, 230.6 kDa; POD-2b, 91.4 kD

MS Intensity: PYC-1 > PCCA-1 > MCCC-1 >> POD-2

CT-TurboID: NEKL-2 80.9 kD; NEKL-3 75.0 kD; MLT-2 115.3 kD; MLT-3 83.5 kD; MLT-4 109.9 kD

#### Experiment 2

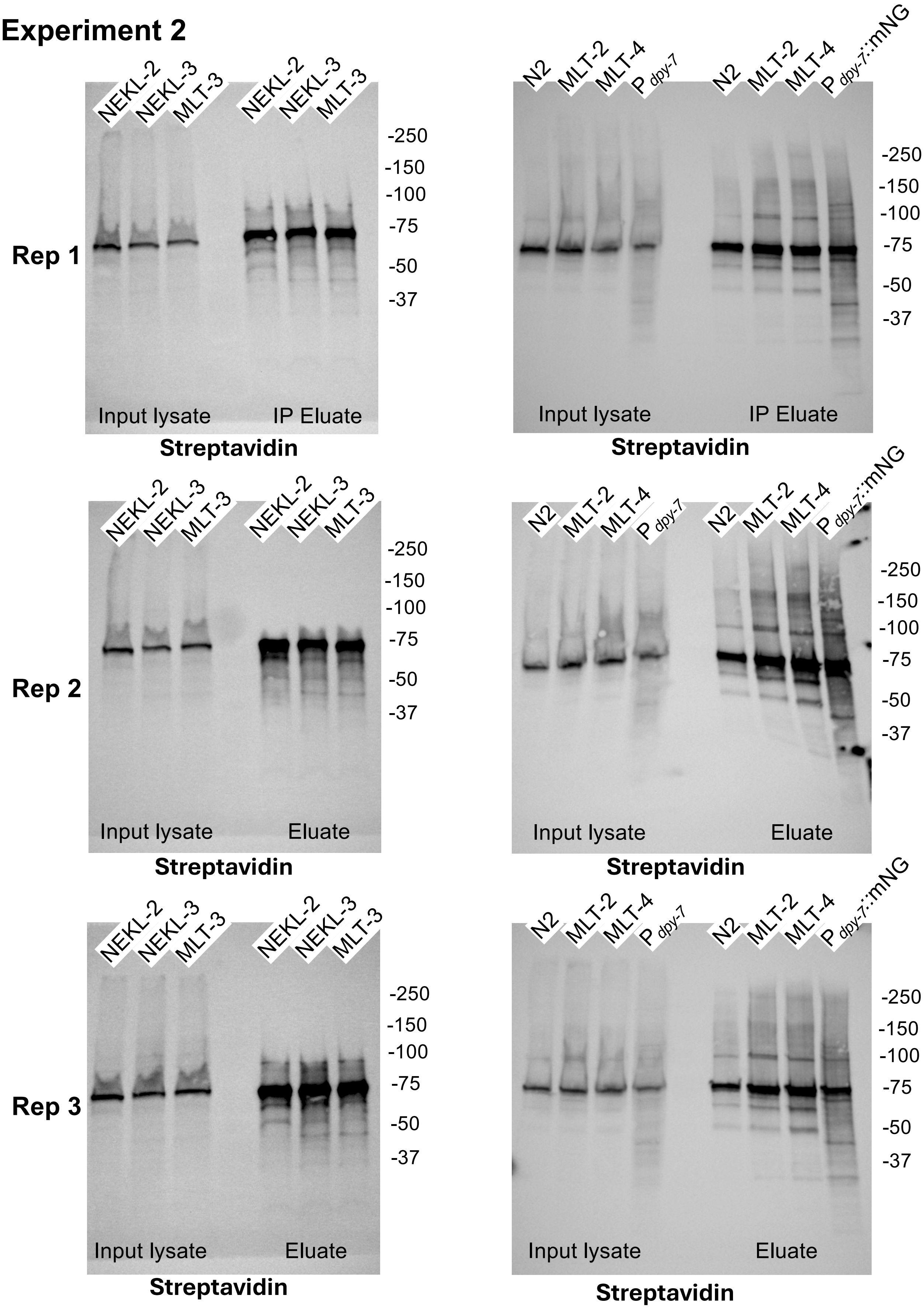


PYC-1a, 129.3 kDa; PYC-1b, 67.8 kDa; PCCA-1, 79.7 kDa; MCCC-1, 73.7 kDa; POD-2a, 230.6 kDa; POD-2b, 91.4 kD

### Experiment 3

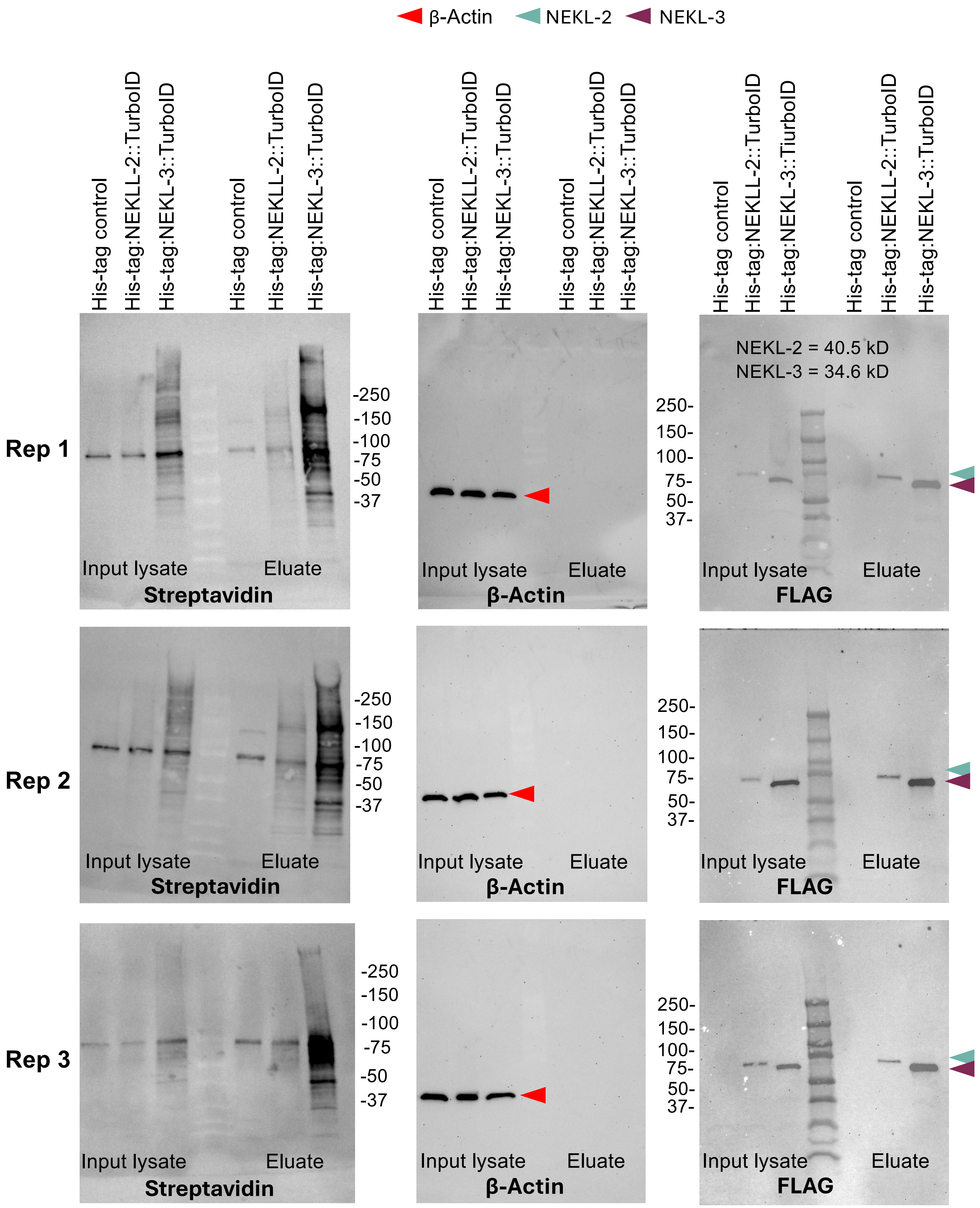


PYC-1a, 129.3 kDa; PYC-1b, 67.8 kDa; PCCA-1, 79.7 kDa; MCCC-1, 73.7 kDa; POD-2a, 230.6 kDa; POD-2b, 91.4 kD

CT-TurboID: NEKL-2 80.9 kDa; NEKL-3 75.0 kDa

### Experiment 4

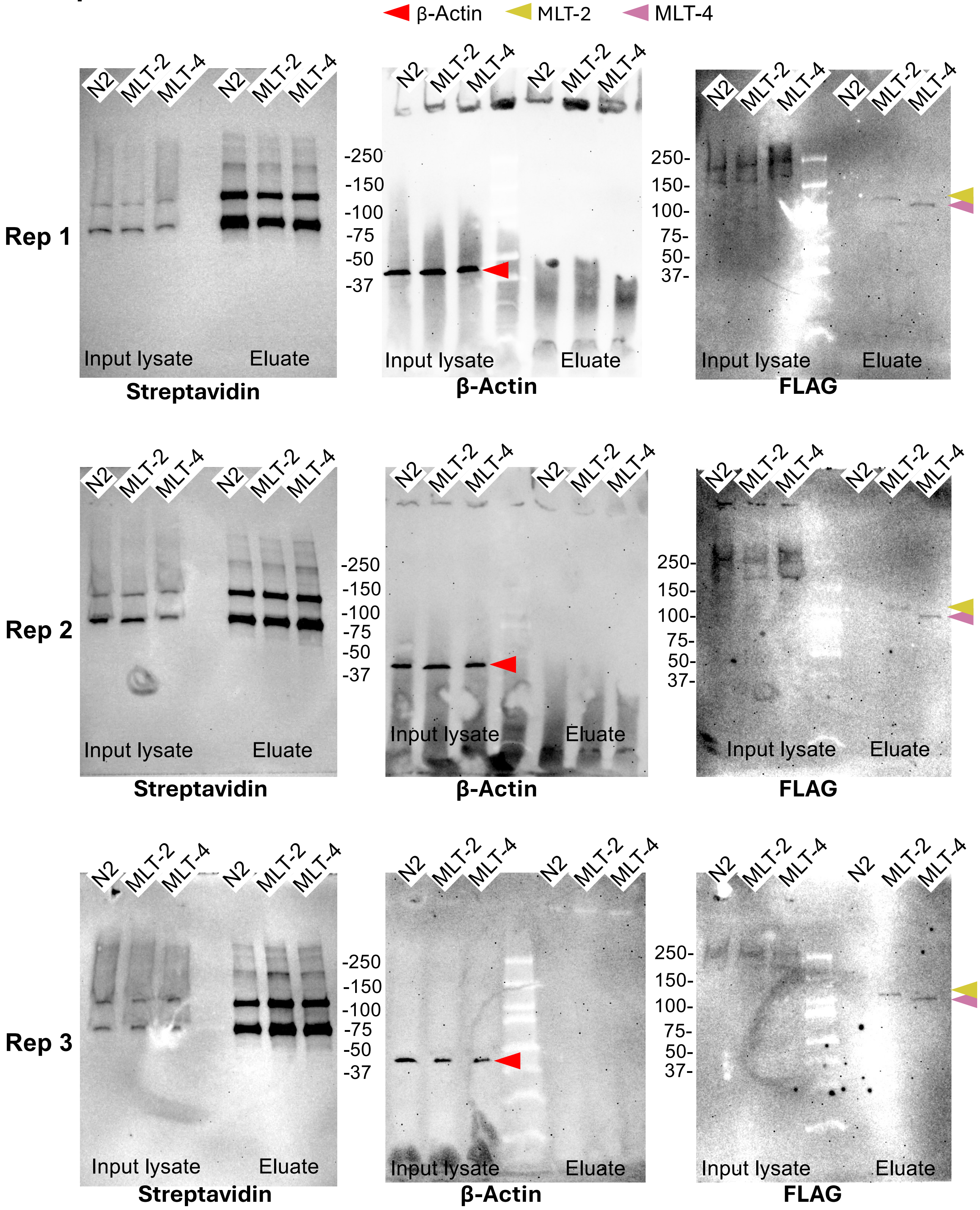


MLT-2 74.9 kDa; MLT-4 69.5 kDa

PYC-1a, 129.3 kDa; PYC-1b, 67.8 kDa; PCCA-1, 79.7 kDa; MCCC-1, 73.7 kDa; POD-2a, 230.6 kDa; POD-2b, 91.4 kD

CT-TurboID: MLT-2 115.3 kDa; MLT-4 109.9 kDa

#### Experiment 5

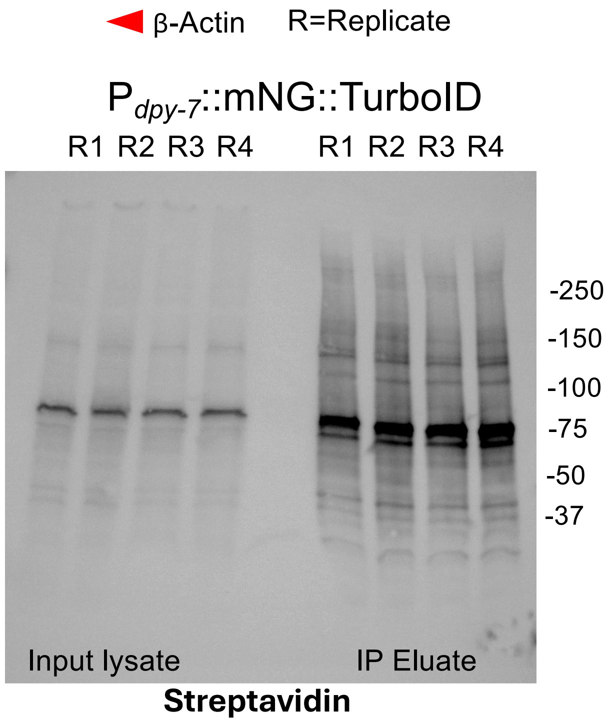


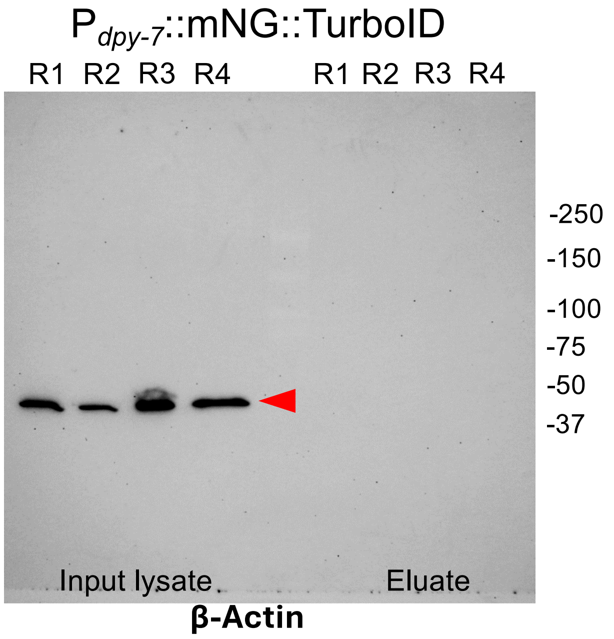


PYC-1a, 129.3 kDa; PYC-1b, 67.8 kDa; PCCA-1, 79.7 kDa; MCCC-1, 73.7 kDa; POD-2a, 230.6 kDa; POD-2b, 91.4 kD

### Experiment 6

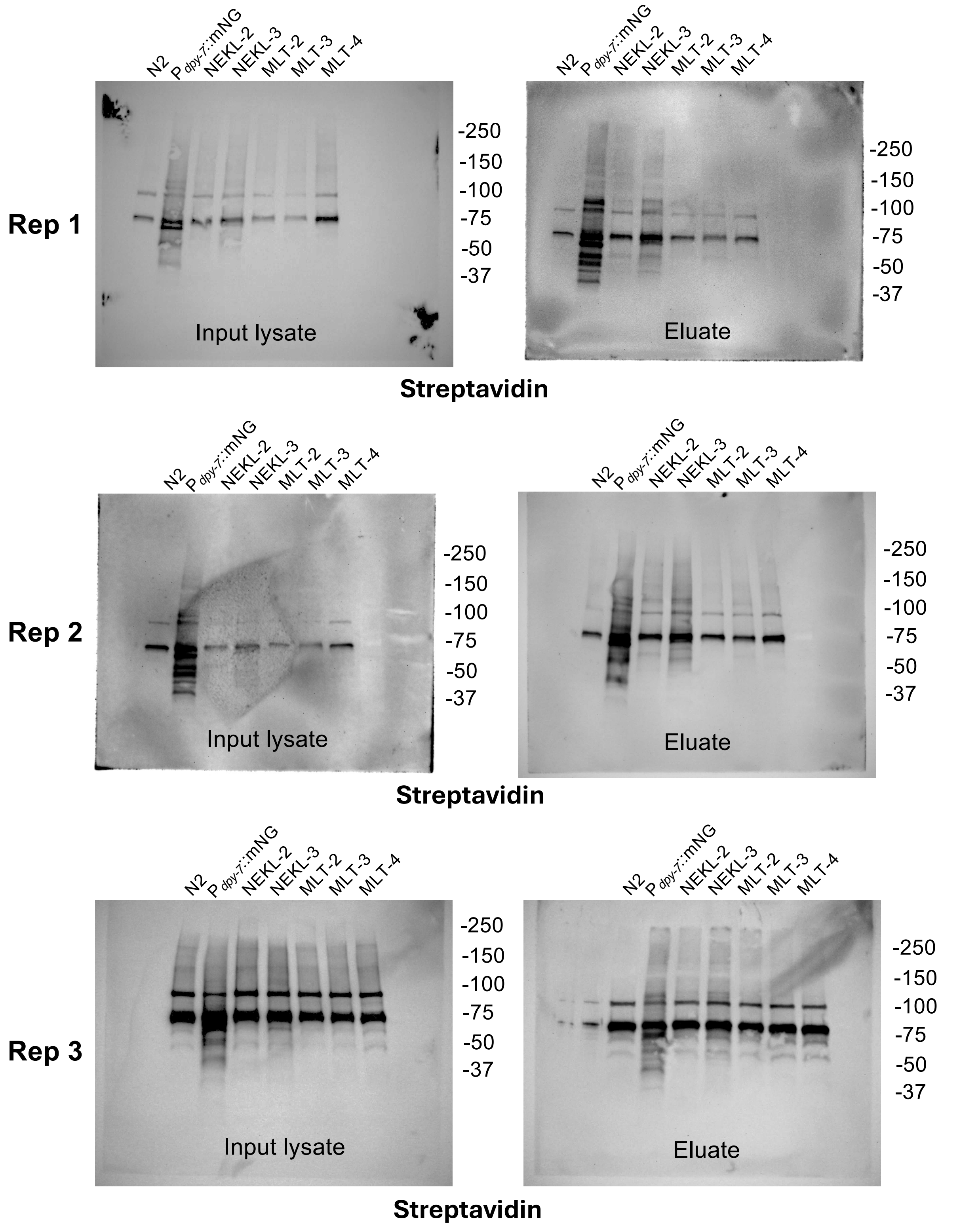


PYC-1a, 129.3 kDa; PYC-1b, 67.8 kDa; PCCA-1, 79.7 kDa; MCCC-1, 73.7 kDa; POD-2a, 230.6 kDa; POD-2b, 91.4 kD

### Experiment 6

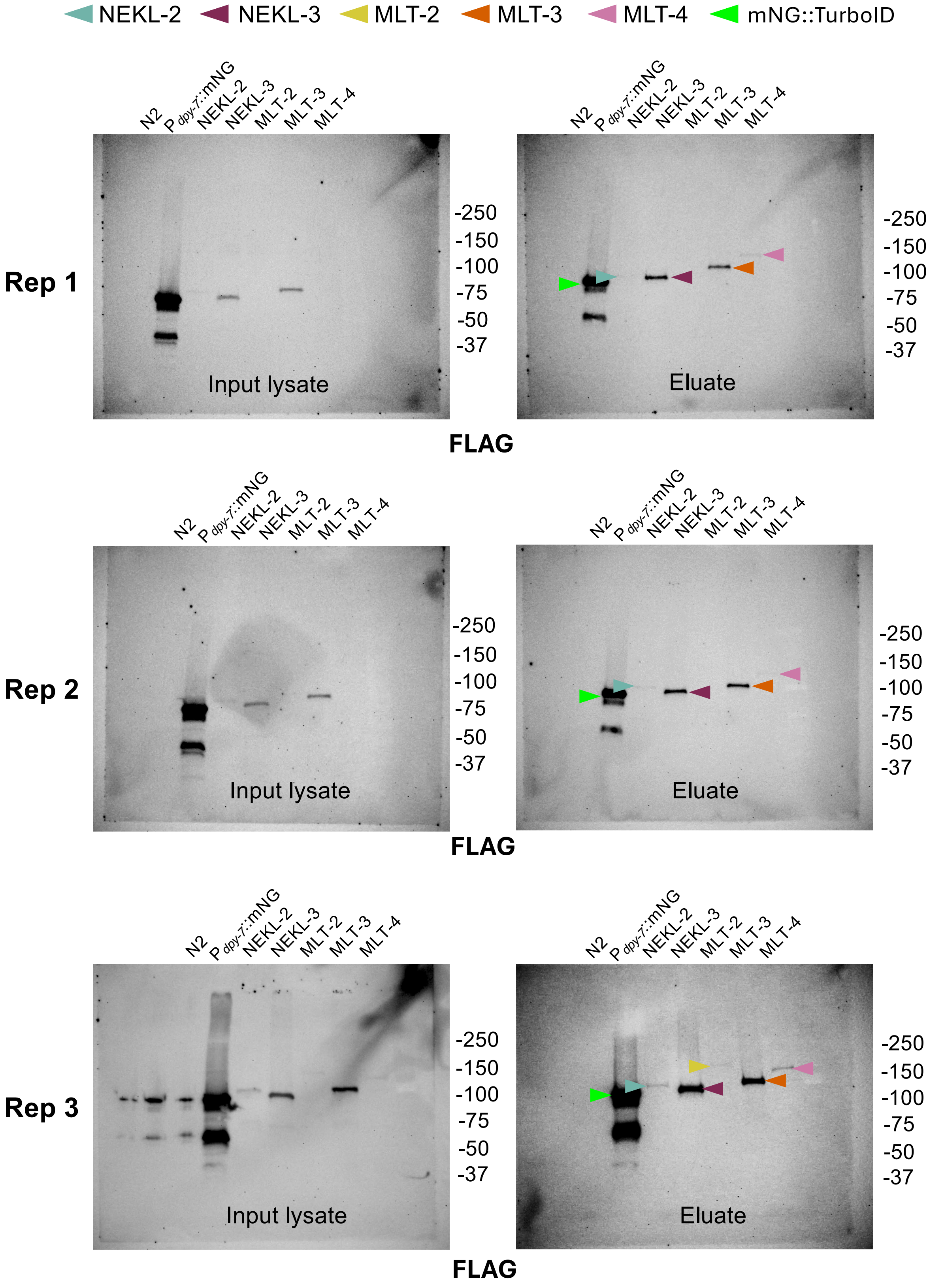


CT-TurboID: NEKL-2 80.9 kDa; NEKL-3 75.0 kDa; MLT-2 115.3 kDa; MLT-3 83.5 kDa; MLT-4 109.9 kDa

#### Experiment 6

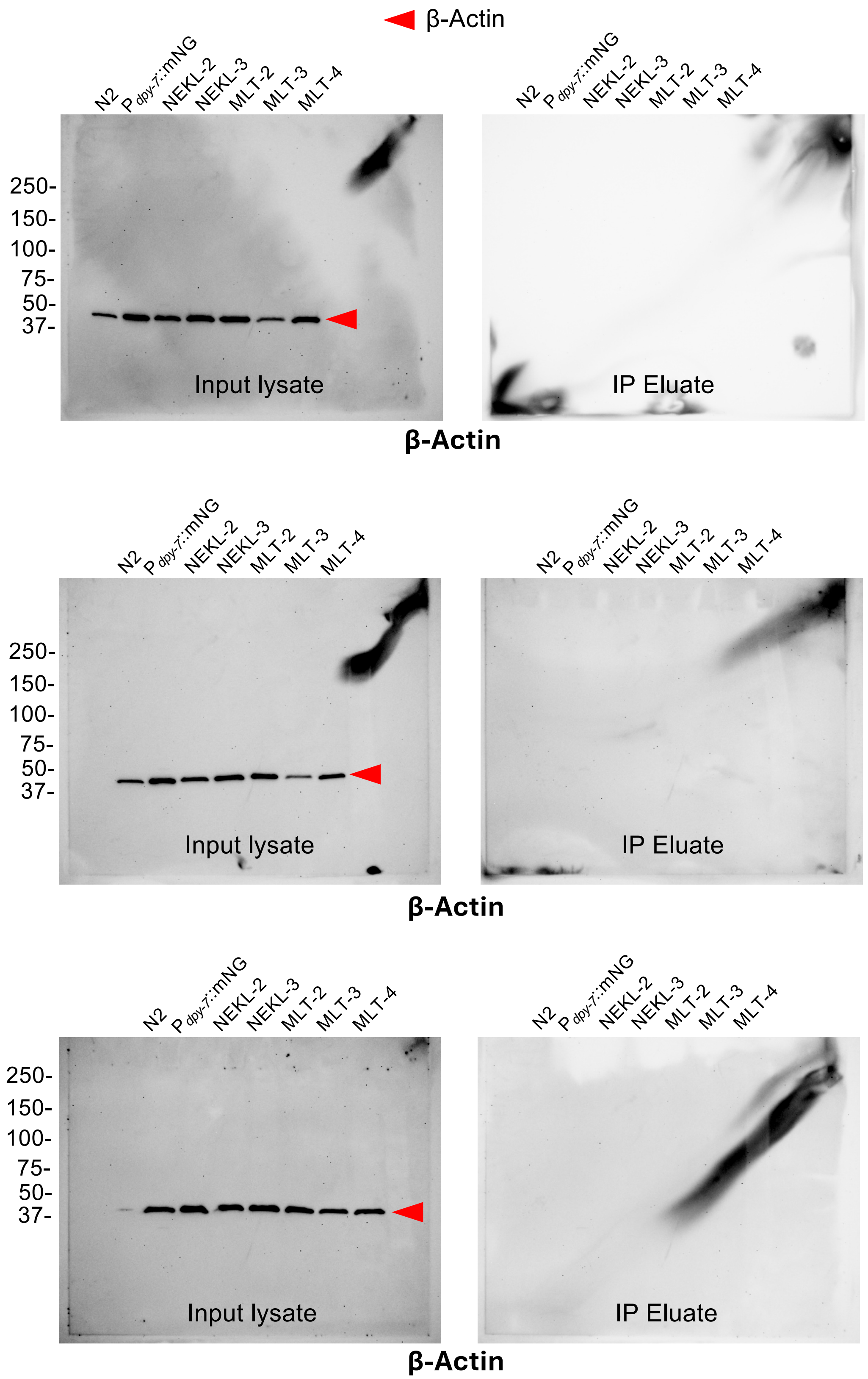

### Experiment 7

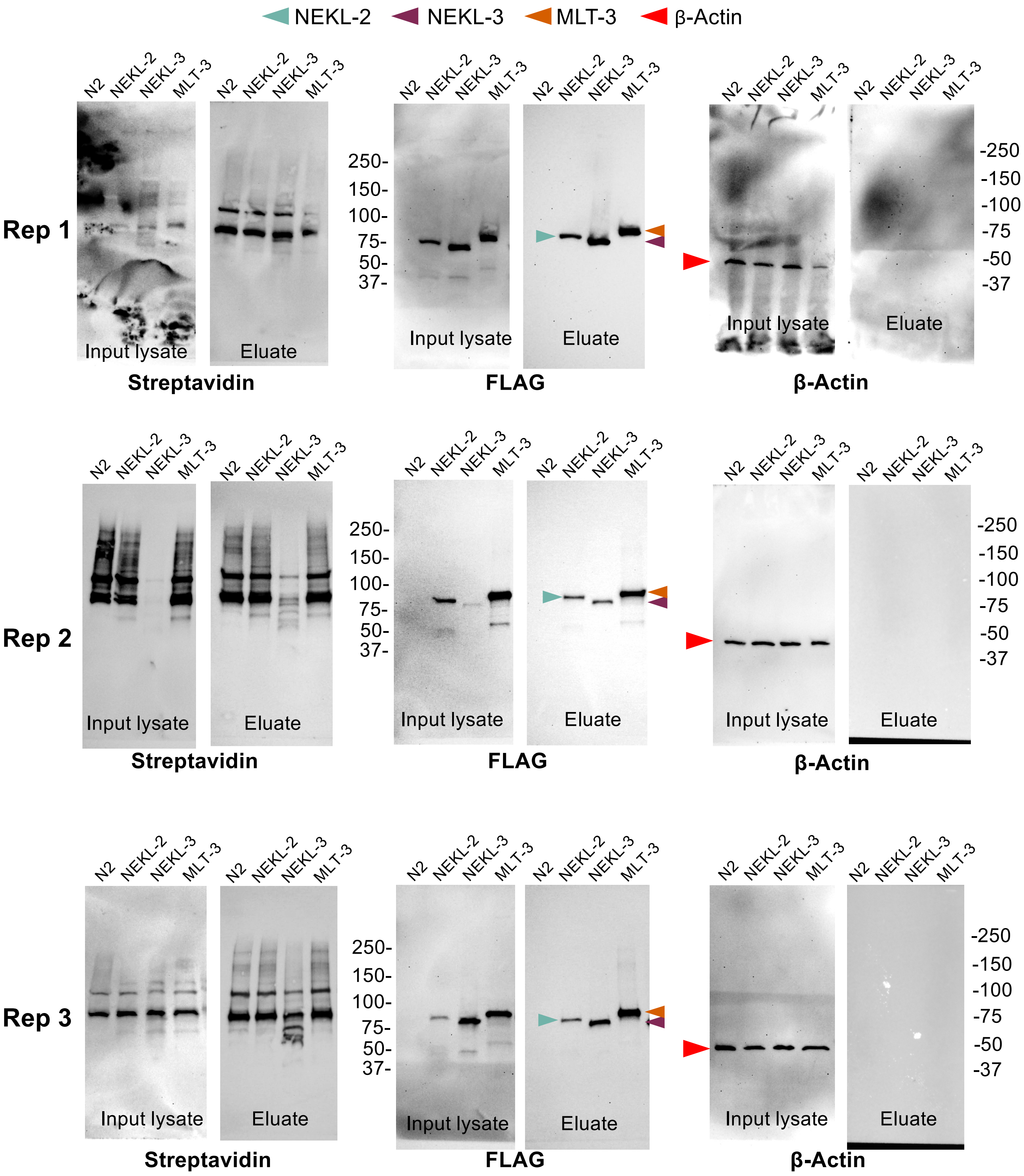


PYC-1a, 129.3 kDa; PYC-1b, 67.8 kDa; PCCA-1, 79.7 kDa; MCCC-1, 73.7 kDa; POD-2a, 230.6 kDa; POD-2b, 91.4 kD

CT-TurboID: NEKL-2 80.9 kDa; NEKL-3 75.0 kDa; MLT-3 83.5 kDa

### Experiment 8

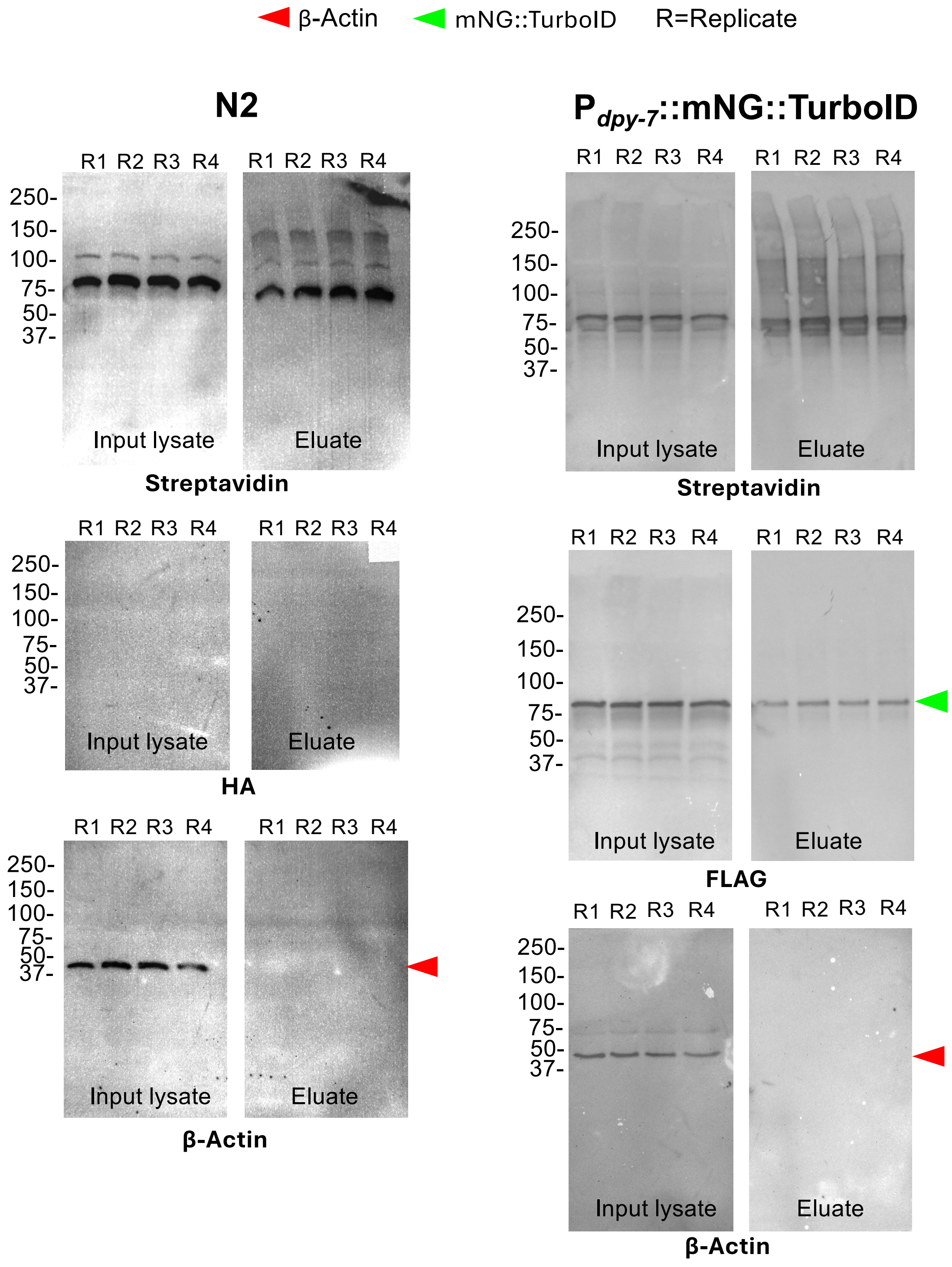


PYC-1a, 129.3 kDa; PYC-1b, 67.8 kDa; PCCA-1, 79.7 kDa; MCCC-1, 73.7 kDa; POD-2a, 230.6 kDa; POD-2b, 91.4 kD

### Experiment 8

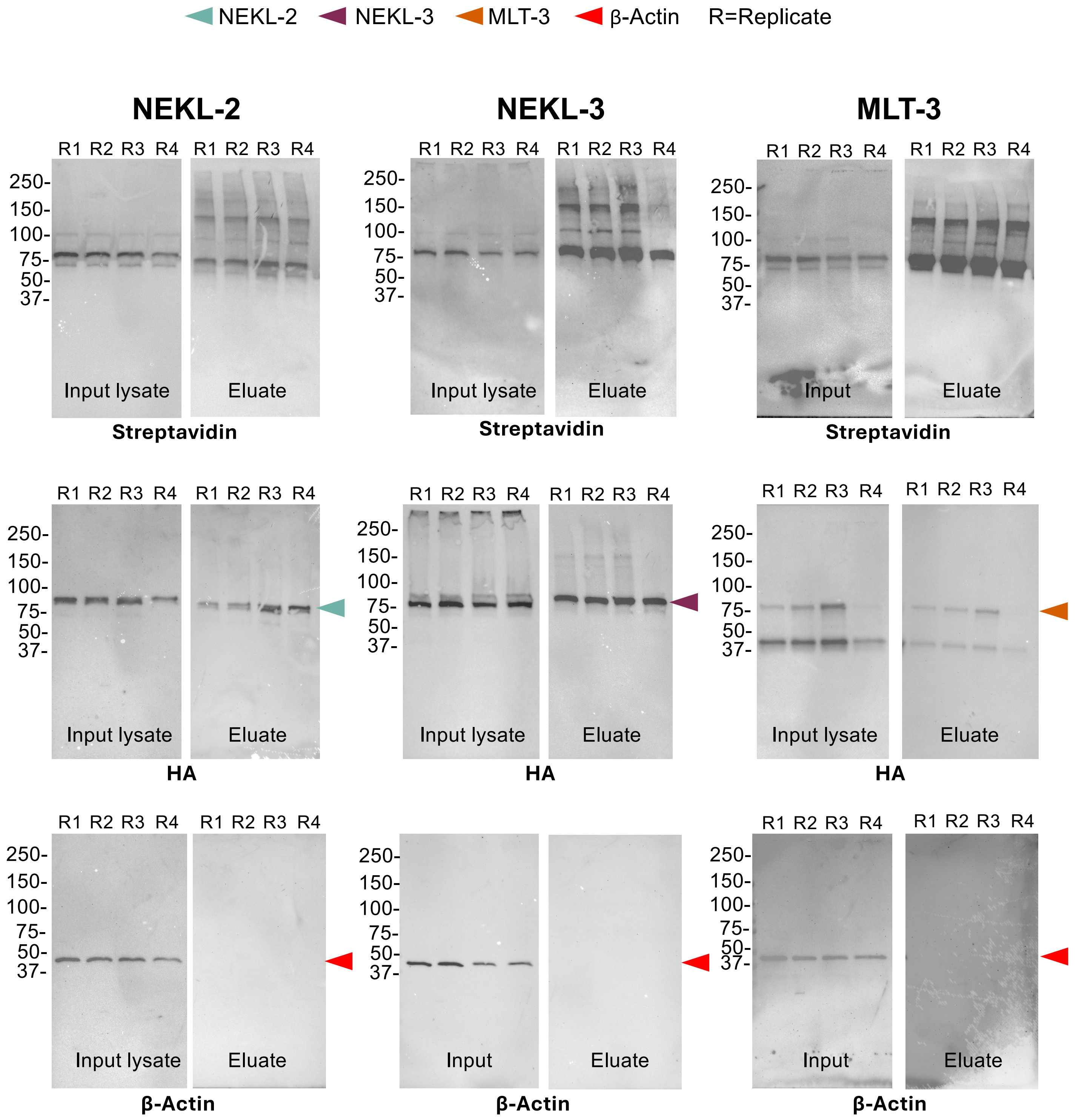


PYC-1a, 129.3 kDa; PYC-1b, 67.8 kDa; PCCA-1, 79.7 kDa; MCCC-1, 73.7 kDa; POD-2a, 230.6 kDa; POD-2b, 91.4 kD

NT-TurboID: NEKL-2 81.0 kDa; NEKL-3 75.1 kDa; MLT-3 83.4 kDa
