## Supplementary material for "Proximity Labeling of NIMA Kinase Complex Components in *C. elegans*": S1 Information

**S2 METHODS**

**C-terminal fusions of TurboID**

Linkers TurboID FLAG-3x **40.4 kD**

**Protein**

GGGGSGSGGRGSGSGSGSGSGSGSGSGKDNTVPLKLIALLANGEFHSGEQLGETLGMSRAAI
NKHIQTLRDWGVDVFTVPGKGYSLPEPIPLLNAKQILGQLDGGSVAVLPVVDSTNQYLLDRIGE
LKSGDACIAEYQQAGRGSRGRKWFSPFGANLYLSMFWRLKRGPAAIGLGPVIGIVMAEALRKLG
ADKVRVKWPNDLYLQDRKLAGILVELAGITGDAAQIVIGAGINVAMRRVEESVVNQGWITLQEA
GINLDRNTLAATLIRELRAALELFEQEGLAPYLPRWEKLDNFINRPVKLIIGDKEIFGISRGID
KQGALLLEQDGVIKPWMGGEISLRSAEKAGGDYKDDDDKRDYKDDDDKRDYKDDDDK

**DNA**

GGAGGTGGTGGATCAGGCTCGGGAGGTCGAGGCTCAGGATCCGGTTCCGGCTCCGGCTCTGGTT
CCGGTTCGGGTTCCGGTTCTGGAAAGGATAACACCGTTCCACTTAAGCTTATCGCCCTTCTTGC
CAACGGAGAATTCCACTCTGGAGAGCAACTTGGAGAGACTCTTGGAATGTCCCGTGCTGCCATC
AACAAGCATATCCAAACCCTTCGTGATTGGGGAGTTGATGTTTTCACTGTTCCAGGAAAGgtaa
gtttaaacatatatatactaactaaccctgattatttaaattttcagGGATACTCCCTTCCAGA
GCCAATCCCACTTCTTAACGCCAAGCAAATCCTTGGACAACTTGATGGAGGATCCGTCGCTGTC
CTTCCAGTTGTTGATTCCACCAACCAATACCTTCTTGACCGTATCGGAGAGCTTAAGTCTGGAG
ACGCCTGCATCGCTGAGTACCAACAAGCTGGACGCGGATCTCGCGGACGCAAGTGGTTCTCCCC
ATTCGGAGCCAACCTTTACCTTTCTATGTTCTGGCGTCTTAAGCGTGGACCAGCTGCTATCGGA
CTTGGACCAGTTATCGGAATCGTTATGGCTGAGGCCCTTCGTAAGCTTGGAGCTGATAAGgtaa
gtttaaacagttcggtactaactaaccatacatatttaaattttcagGTTCGTGTTAAGTGGCC
AAACGATCTTTACCTTCAAGACCGTAAGCTTGCTGGAATCCTTGTCGAGCTTGCTGGAATCACC
GGAGACGCCGCTCAAATCGTTATCGGAGCTGGAATCAACGTTGCCATGCGTCGTGTTGAGGAGT
CTGTTGTTAACCAAGGATGGATCACTCTTCAAGAGGCTGGAATCAACCTTGATCGTAACACCCT
TGCTGCCACCCTTATCCGTGAGCTTCGTGCTGCCCTTGAGCTTTTCGAGCAAGAGGGACTTGCC
CCATACCTTCCACGCTGGGAGAAGCTTGACAACTTCATCAACCGCCCAGTTAAGCTTATCATCG
GAGATAAGGAAATCTTCGGAATCTCTCGCGGAATCGACAAGCAAGGAGCTCTTCTTCTTGAGCA
AGATGGAGTCATTAAGCCATGGATGGGAGGAGAGATTTCCCTTCGTTCCGCTGAGAAGGCCGGA
GGAGATTATAAAGACGATGACGATAAGCGTGACTACAAGGACGACGACGACAAGCGTGATTACA
AGGATGACGATGACAAG

Lower case in TurboID sequence indicates synthetic introns.

Final coding codon of NEKL-MLT bait proteins directly preceeds the N-terminal linker sequence.

Endogenous stop codon of NEKL-MLT bait proteins directly follows the C-terminal FLAG-3x sequence.

**N-terminal fusions of TurboID**

Linkers TurboID HA-3x **40.5 kD**

**Protein**

PYDVPDYAGSYPYDVPDYAGSYPYDVPDYAGGGKDNTVPLKLIALLANGEFHSGEQLGETLGMS
RAAINKHIQTLRDWGVDVFTVPGKGYSLPEPIPLLNAKQILGQLDGGSVAVLPVVDSTNQYLLD
RIGELKSGDACIAEYQQAGRGSRGRKWFSPFGANLYLSMFWRLKRGPAAIGLGPVIGIVMAEAL
RKLGADKVRVKWPNDLYLQDRKLAGILVELAGITGDAAQIVIGAGINVAMRRVEESVVNQGWIT
LQEAGINLDRNTLAATLIRELRAALELFEQEGLAPYLPRWEKLDNFINRPVKLIIGDKEIFGIS
RGIDKQGALLLEQDGVIKPWMGGEISLRSAEKGGGGSGSGGRGSGSGSGSGSGSGSGSG

**DNA**

TACCCATATGACGTTCCAGACTACGCGGGTAGTTATCCGTACGACGTTCCGGATTACGCTGGAT
CTTACCCTTACGACGTACCTGACTACGCTGGAGGTGGTAAGGATAACACCGTTCCACTTAAGCT
TATCGCCCTTCTTGCCAACGGAGAATTCCACTCTGGAGAGCAACTTGGAGAGACTCTTGGAATG
TCCCGTGCTGCCATCAACAAGCATATCCAAACCCTTCGTGATTGGGGAGTTGATGTTTTCACTG
TTCCAGGAAAGgtaagtttaaacatatatatactaactaaccctgattatttaaattttcagGG
ATACTCCCTTCCAGAGCCAATCCCACTTCTTAACGCCAAGCAAATCCTTGGACAACTTGATGGA
GGATCCGTCGCTGTCCTTCCAGTTGTTGATTCCACCAACCAATACCTTCTTGACCGTATCGGAG
AGCTTAAGTCTGGAGACGCCTGCATCGCTGAGTACCAACAAGCTGGACGCGGATCTCGCGGACG
CAAGTGGTTCTCCCCATTCGGAGCCAACCTTTACCTTTCTATGTTCTGGCGTCTTAAGCGTGGA
CCAGCTGCTATCGGACTTGGACCAGTTATCGGAATCGTTATGGCTGAGGCCCTTCGTAAGCTTG
GAGCTGATAAGgtaagtttaaacagttcggtactaactaaccatacatatttaaattttcagGT
TCGTGTTAAGTGGCCAAACGATCTTTACCTTCAAGACCGTAAGCTTGCTGGAATCCTTGTCGAG
CTTGCTGGAATCACCGGAGACGCCGCTCAAATCGTTATCGGAGCTGGAATCAACGTTGCCATGC
GTCGTGTTGAGGAGTCTGTTGTTAACCAAGGATGGATCACTCTTCAAGAGGCTGGAATCAACCT
TGATCGTAACACCCTTGCTGCCACCCTTATCCGTGAGCTTCGTGCTGCCCTTGAGCTTTTCGAG
CAAGAGGGACTTGCCCCATACCTTCCACGCTGGGAGAAGCTTGACAACTTCATCAACCGCCCAG
TTAAGCTTATCATCGGAGATAAGGAAATCTTCGGAATCTCTCGCGGAATCGACAAGCAAGGAGC
TCTTCTTCTTGAGCAAGATGGAGTCATTAAGCCATGGATGGGAGGAGAGATTTCCCTTCGTTCC
GCTGAGAAGGGAGGTGGTGGATCAGGCTCGGGAGGTCGAGGCTCAGGATCCGGTTCCGGCTCCG
GCTCTGGTTCCGGTTCGGGTTCCGGTTCTGGA

Lower case in TurboID sequence indicates synthetic introns.

First codon of NEKL-MLT bait proteins (ATG) directly precedes the HA-3x tag sequence.

Second codon of NEKL-MLT bait proteins directly follows the C-terminal linker sequence.
