## Supplementary material for "Proximity Labeling of NIMA Kinase Complex Components in *C. elegans*": S2 Information

**Exp 1 FASP Methods – Orbitrap Exploris DIA**

Protein samples were reduced, alkylated, and digested on-bead using filter-aided sample preparation [*Nature Methods* **6:** 359-62 (2009)] with sequencing grade modified porcine trypsin (Promega).

Tryptic peptides were trapped and eluted on 3.5um CSH C18 resin (Waters) (4mm x 75um) then separated by reverse phase XSelect CSH C18 2.5 um resin (Waters) on an in-line 150 x 0.075 mm column using an UltiMate 3000 RSLCnano system (Thermo). Peptides were eluted at a flow rate of 0.300uL/min using a 60 min gradient from 98% Buffer A:2% Buffer B to 95:5 at 2.0 minutes to 80:20 at 39.0 minutes to 60:40 at 48.0 minutes to 10:90 at 49.0 minutes and hold until 53.0 minutes and then equilibrated back to 98:2 at 53.1 minutes until 60 minutes. Eluted peptides were ionized by electrospray (2.4 kV) through a heated capillary (275°C) followed by data collection on an Orbitrap Exploris 480 mass spectrometer (Thermo Scientific).

Precursor spectra were acquired with a scan from 385-1015 Th at a resolution set to 60,000 with 100% AGC, max time of 50 msec, and an RF parameter at 40%. DIA was configured on the Orbitrap 480 to acquire 50 x 12 Th isolation windows at 15,000 resolution, normalized AGC target 500%, maximum injection time 40 ms). A second DIA was acquired in a staggered window (12 Th) pattern with optimized window placements.

Buffer A = 0.1% formic acid, 0.5% acetonitrile
Buffer B = 0.1% formic acid, 99.9% acetonitrile

Following data acquisition, data were searched using Spectronaut (Biognosys version 18.3) against the UniProt *Caenorhabditis elegans* database (August 2023) using the directDIA method with an identification precursor and protein q-value cutoff of 1%, generate decoys set to true, the protein inference workflow set to maxLFQ, inference algorithm set to IDPicker, quantity level set to MS2, cross-run normalization set to false, and the protein grouping quantification set to median peptide and precursor quantity. Protein MS2 intensity values were assessed for quality using ProteiNorm (Graw *et al.*). The data was normalized using VSN (Huber et al) and analyzed using proteoDA to perform statistical analysis using Linear Models for Microarray Data (limma) with empirical Bayes (eBayes) smoothing to the standard errors (Thurman et al, Ritchie et al). Proteins with an FDR adjusted p-value < 0.05 and a fold change > 2 were considered significant.

**Acknowledgement:**
IDeA National Resource for Quantitative Proteomics and NIH/NIGMS grant R24GM137786.

**Exp 2 FASP Methods – Orbitrap Exploris DIA**

Protein samples were reduced, alkylated, and digested on-bead using filter-aided sample preparation [*Nature Methods* **6:** 359-62 (2009)] with sequencing grade modified porcine trypsin (Promega). Tryptic peptides were then separated by reverse phase XSelect CSH C18 2.5 um resin (Waters) on an in-line 150 x 0.075 mm column using an UltiMate 3000 RSLCnano system (Thermo). Peptides were eluted using a 60 min gradient from 98:2 to 65:35 buffer A:B ratio. Eluted peptides were ionized by electrospray (2.2 kV) followed by mass spectrometric analysis on an Orbitrap Exploris 480 mass spectrometer (Thermo). To assemble a chromatogram library, six gas-phase fractions were acquired on the Orbitrap Exploris with 4 m/z DIA spectra (4 m/z precursor isolation windows at 30,000 resolution, normalized AGC target 100%, maximum inject time 66 ms) using a staggered window pattern from narrow mass ranges using optimized window placements. Precursor spectra were acquired after each DIA duty cycle, spanning the m/z range of the gas-phase fraction (i.e. 496-602 m/z, 60,000 resolution, normalized AGC target 100%, maximum injection time 50 ms). For wide-window acquisitions, the Orbitrap Exploris was configured to acquire a precursor scan (385-1015 m/z, 60,000 resolution, normalized AGC target 100%, maximum injection time 50 ms) followed by 50x 12 m/z DIA spectra (12 m/z precursor isolation windows at 15,000 resolution, normalized AGC target 100%, maximum injection time 33 ms) using a staggered window pattern with optimized window placements. Precursor spectra were acquired after each DIA duty cycle.

Buffer A = 0.1% formic acid, 0.5% acetonitrile
Buffer B = 0.1% formic acid, 99.9% acetonitrile

Following data acquisition, data were searched using Spectronaut (Biognosys version 18.5) against the UniProt *Caenorhabditis elegans* database (April 2023) using the directDIA method with an identification precursor and protein q-value cutoff of 1%, generate decoys set to true, the protein inference workflow set to maxLFQ, inference algorithm set to IDPicker, quantity level set to MS2, cross-run normalization set to false, and the protein grouping quantification set to median peptide and precursor quantity. Protein MS2 intensity values were assessed for quality using ProteiNorm (Graw *et al.*). The data was normalized using VSN (Huber et al) and analyzed using proteoDA to perform statistical analysis using Linear Models for Microarray Data (limma) with empirical Bayes (eBayes) smoothing to the standard errors (Thurman et al, Ritchie et al). Proteins with an FDR adjusted p-value < 0.05 and a fold change > 2 were considered significant.

**Acknowledgement:**
IDeA National Resource for Quantitative Proteomics and NIH/NIGMS grant R24GM137786.

**Exp 3 FASP Methods – Orbitrap Exploris DIA**

Protein samples were reduced, alkylated, and digested on-bead using filter-aided sample preparation [*Nature Methods* **6:** 359-62 (2009)] with sequencing grade modified porcine trypsin (Promega). Tryptic peptides were then separated by reverse phase XSelect CSH C18 2.5 um resin (Waters) on an in-line 150 x 0.075 mm column using an UltiMate 3000 RSLCnano system (Thermo). Peptides were eluted using a 60 min gradient from 98:2 to 65:35 buffer A:B ratio. Eluted peptides were ionized by electrospray (2.2 kV) followed by mass spectrometric analysis on an Orbitrap Exploris 480 mass spectrometer (Thermo). To assemble a chromatogram library, six gas-phase fractions were acquired on the Orbitrap Exploris with 4 m/z DIA spectra (4 m/z precursor isolation windows at 30,000 resolution, normalized AGC target 100%, maximum inject time 66 ms) using a staggered window pattern from narrow mass ranges using optimized window placements. Precursor spectra were acquired after each DIA duty cycle, spanning the m/z range of the gas-phase fraction (i.e. 496-602 m/z, 60,000 resolution, normalized AGC target 100%, maximum injection time 50 ms). For wide-window acquisitions, the Orbitrap Exploris was configured to acquire a precursor scan (385-1015 m/z, 60,000 resolution, normalized AGC target 100%, maximum injection time 50 ms) followed by 50x 12 m/z DIA spectra (12 m/z precursor isolation windows at 15,000 resolution, normalized AGC target 100%, maximum injection time 33 ms) using a staggered window pattern with optimized window placements. Precursor spectra were acquired after each DIA duty cycle.

Buffer A = 0.1% formic acid, 0.5% acetonitrile
Buffer B = 0.1% formic acid, 99.9% acetonitrile

Following data acquisition, data were searched using Spectronaut (Biognosys version 18.5) against the UniProt *Caenorhabditis elegans* database (September 2023) using the directDIA method with an identification precursor and protein q-value cutoff of 1%, generate decoys set to true, the protein inference workflow set to maxLFQ, inference algorithm set to IDPicker, quantity level set to MS2, cross-run normalization set to false, and the protein grouping quantification set to median peptide and precursor quantity. Protein MS2 intensity values were assessed for quality using ProteiNorm (Graw *et al.*). The data was normalized using VSN (Huber et al) and analyzed using proteoDA to perform statistical analysis using Linear Models for Microarray Data (limma) with empirical Bayes (eBayes) smoothing to the standard errors (Thurman et al, Ritchie et al). Proteins with an FDR adjusted p-value < 0.05 and a fold change > 2 were considered significant.

**Acknowledgement:**
IDeA National Resource for Quantitative Proteomics and NIH/NIGMS grant R24GM137786.

**Exp 4 FASP Methods – Orbitrap Exploris DIA**

Protein samples were reduced, alkylated, and digested on-bead using filter-aided sample preparation [*Nature Methods* **6:** 359-62 (2009)] with sequencing grade modified porcine trypsin (Promega). Tryptic peptides were then separated by reverse phase XSelect CSH C18 2.5 um resin (Waters) on an in-line 150 x 0.075 mm column using an UltiMate 3000 RSLCnano system (Thermo). Peptides were eluted using a 60 min gradient from 98:2 to 65:35 buffer A:B ratio. Eluted peptides were ionized by electrospray (2.2 kV) followed by mass spectrometric analysis on an Orbitrap Exploris 480 mass spectrometer (Thermo). To assemble a chromatogram library, six gas-phase fractions were acquired on the Orbitrap Exploris with 4 m/z DIA spectra (4 m/z precursor isolation windows at 30,000 resolution, normalized AGC target 100%, maximum inject time 66 ms) using a staggered window pattern from narrow mass ranges using optimized window placements. Precursor spectra were acquired after each DIA duty cycle, spanning the m/z range of the gas-phase fraction (i.e. 496-602 m/z, 60,000 resolution, normalized AGC target 100%, maximum injection time 50 ms). For wide-window acquisitions, the Orbitrap Exploris was configured to acquire a precursor scan (385-1015 m/z, 60,000 resolution, normalized AGC target 100%, maximum injection time 50 ms) followed by 50x 12 m/z DIA spectra (12 m/z precursor isolation windows at 15,000 resolution, normalized AGC target 100%, maximum injection time 33 ms) using a staggered window pattern with optimized window placements. Precursor spectra were acquired after each DIA duty cycle.

Buffer A = 0.1% formic acid, 0.5% acetonitrile
Buffer B = 0.1% formic acid, 99.9% acetonitrile

Following data acquisition, data were searched using Spectronaut (Biognosys version 18.5) against the UniProt *Caenorhabditis elegans* database (September 2023) using the directDIA method with an identification precursor and protein q-value cutoff of 1%, generate decoys set to true, the protein inference workflow set to maxLFQ, inference algorithm set to IDPicker, quantity level set to MS2, cross-run normalization set to false, and the protein grouping quantification set to median peptide and precursor quantity. Protein MS2 intensity values were assessed for quality using ProteiNorm (Graw *et al.*). The data was normalized using VSN (Huber et al) and analyzed using proteoDA to perform statistical analysis using Linear Models for Microarray Data (limma) with empirical Bayes (eBayes) smoothing to the standard errors (Thurman et al, Ritchie et al). Proteins with an FDR adjusted p-value < 0.05 and a fold change > 2 were considered significant.

**Acknowledgement:**
IDeA National Resource for Quantitative Proteomics and NIH/NIGMS grant R24GM137786.

Buffer A = 0.1% formic acid, 0.5% acetonitrile
Buffer B = 0.1% formic acid, 99.9% acetonitrile

**Data Analysis**

Proteins were identified and quantified by searching the UniprotKB database restricted to *Caenorhabditis elegans* (February 2024) using MaxQuant (version 2.2.0.0, Max Planck Institute). The database search parameters included selecting the MS1 reporter type, trypsin digestion with up to two missed cleavages, fixed modifications for carbamidomethyl of cysteine, variable modifications for oxidation on methionine and acetyl on N-terminus, the precursor ion tolerance of 5 ppm for the first search and 3 ppm for the main search, and label-free quantitation with iBAQ. Peptide and protein identifications were accepted using the 1.0% false discovery rate identification threshold. Protein probabilities were assigned by the Protein Prophet algorithm (Nesvizhskii et al 2003).

MaxQuant iBAQ intensities for each sample were assessed for quality and differential abundance using proteoDA (Thurman et al. 2023; Graw et al. 2020). The data was normalized using VSN (Huber et al. 2002) and statistical analysis was performed using linear models for microarray data (limma) with empirical Bayes (eBayes) smoothing to the standard errors (Ritchie et al.2015). Proteins with an FDR adjusted p-value < 0.05 and a fold change > 2 were considered significant.

**Acknowledgement:**
IDeA National Resource for Quantitative Proteomics and NIH/NIGMS grant R24GM137786.

### Exp 6 FASP Methods – Orbitrap Exploris DIA

Protein samples were reduced, alkylated, and digested on-bead using filter-aided sample preparation [*Nature Methods* **6:** 359-62 (2009)] with sequencing grade modified porcine trypsin (Promega).

Tryptic peptides were trapped and eluted on 3.5um CSH C18 resin (Waters) (4mm x 75um) then separated by reverse phase XSelect CSH C18 2.5 um resin (Waters) on an in-line 150 x 0.075 mm column using an UltiMate 3000 RSLCnano system (Thermo). Peptides were eluted at a flow rate of 0.300uL/min using a 60 min gradient from 98% Buffer A:2% Buffer B to 95:5 at 2.0 minutes to 80:20 at 39.0 minutes to 60:40 at 48.0 minutes to 10:90 at 49.0 minutes and hold until 53.0 minutes and then equilibrated back to 98:2 at 53.1 minutes until 60 minutes. Eluted peptides were ionized by electrospray (2.4 kV) through a heated capillary (275°C) followed by data collection on an Orbitrap Exploris 480 mass spectrometer (Thermo Scientific).

Precursor spectra were acquired with a scan from 385-1015 Th at a resolution set to 60,000 with 100% AGC, max time of 50 msec, and an RF parameter at 40%. DIA was configured on the Orbitrap 480 to acquire 50 x 12 Th isolation windows at 15,000 resolution, normalized AGC target 500%, maximum injection time 40 ms). A second DIA was acquired in a staggered window (12 Th) pattern with optimized window placements.

Buffer A = 0.1% formic acid, 0.5% acetonitrile
Buffer B = 0.1% formic acid, 99.9% acetonitrile

Following data acquisition, data were searched using Spectronaut (Biognosys version 19.1) against the UniProt *Caenorhabditis elegans* database (April 2024) using the directDIA method with an identification precursor and protein q-value cutoff of 1%, generate decoys set to true, the protein inference workflow set to maxLFQ, inference algorithm set to IDPicker, quantity level set to MS2, cross-run normalization set to false, and the protein grouping quantification set to median peptide and precursor quantity. Protein MS2 intensity values were assessed for quality using ProteiNorm (Graw *et al.*). The data was normalized using VSN (Huber et al) and analyzed using proteoDA to perform statistical analysis using Linear Models for Microarray Data (limma) with empirical Bayes (eBayes) smoothing to the standard errors (Thurman et al, Ritchie et al). Proteins with an FDR adjusted p-value < 0.05 and a fold change > 2 were considered significant.

**Acknowledgement:**
IDeA National Resource for Quantitative Proteomics and NIH/NIGMS grant R24GM137786.

Tryptic peptides were trapped and eluted on 3.5um CSH C18 resin (Waters) (4mm x 75um) then separated by reverse phase XSelect CSH C18 2.5 um resin (Waters) on an in-line 150 x 0.075 mm column using an UltiMate 3000 RSLCnano system (Thermo). Peptides were eluted at a flow rate of 0.300uL/min using a 60 min gradient from 98% Buffer A:2% Buffer B to 95:5 at 2.0 minutes to 80:20 at 39.0 minutes to 60:40 at 48.0 minutes to 10:90 at 49.0 minutes and hold until 53.0 minutes and then equilibrated back to 98:2 at 53.1 minutes until 60 minutes. Eluted peptides were ionized by electrospray (2.4 kV) through a heated capillary (275°C) followed by data collection on an Orbitrap Exploris 480 mass spectrometer (Thermo Scientific).

Precursor spectra were acquired with a scan from 385-1015 Th at a resolution set to 60,000 with 100% AGC, max time of 50 msec, and an RF parameter at 40%. DIA was configured on the Orbitrap 480 to acquire 50 x 12 Th isolation windows at 15,000 resolution, normalized AGC target 500%, maximum injection time 40 ms). A second DIA was acquired in a staggered window (12 Th) pattern with optimized window placements.

Buffer A = 0.1% formic acid, 0.5% acetonitrile
Buffer B = 0.1% formic acid, 99.9% acetonitrile

Following data acquisition, data were searched using Spectronaut (Biognosys version 19.1) against the UniProt *Caenorhabditis elegans* database (April 2024) using the directDIA method with an identification precursor and protein q-value cutoff of 1%, generate decoys set to true, the protein inference workflow set to maxLFQ, inference algorithm set to IDPicker, quantity level set to MS2, cross-run normalization set to false, and the protein grouping quantification set to median peptide and precursor quantity. Protein MS2 intensity values were assessed for quality using ProteiNorm (Graw *et al.*). The data was normalized using VSN (Huber et al) and analyzed using proteoDA to perform statistical analysis using Linear Models for Microarray Data (limma) with empirical Bayes (eBayes) smoothing to the standard errors (Thurman et al, Ritchie et al). Proteins with an FDR adjusted p-value < 0.05 and a fold change > 2 were considered significant.

**Acknowledgement:**
IDeA National Resource for Quantitative Proteomics and NIH/NIGMS grant R24GM137786.

Tryptic peptides were trapped and eluted on 3.5um CSH C18 resin (Waters) (4mm x 75um) then separated by reverse phase XSelect CSH C18 2.5 um resin (Waters) on an in-line 150 x 0.075 mm column using an UltiMate 3000 RSLCnano system (Thermo). Peptides were eluted at a flow rate of 0.300uL/min using a 60 min gradient from 98% Buffer A:2% Buffer B to 95:5 at 2.0 minutes to 80:20 at 39.0 minutes to 60:40 at 48.0 minutes to 10:90 at 49.0 minutes and hold until 53.0 minutes and then equilibrated back to 98:2 at 53.1 minutes until 60 minutes. Eluted peptides were ionized by electrospray (2.4 kV) through a heated capillary (275°C) followed by data collection on an Orbitrap Exploris 480 mass spectrometer (Thermo Scientific).

Precursor spectra were acquired with a scan from 385-1015 Th at a resolution set to 60,000 with 100% AGC, max time of 50 msec, and an RF parameter at 40%. DIA was configured on the Orbitrap 480 to acquire 50 x 12 Th isolation windows at 15,000 resolution, normalized AGC target 500%, maximum injection time 40 ms). A second DIA was acquired in a staggered window (12 Th) pattern with optimized window placements.

Buffer A = 0.1% formic acid, 0.5% acetonitrile
Buffer B = 0.1% formic acid, 99.9% acetonitrile

Following data acquisition, data were searched using Spectronaut (Biognosys version 19.5) against the UniProt *Caenorhabditis elegans* database (June 2024) using the directDIA method with an identification precursor and protein q-value cutoff of 1%, generate decoys set to true, the protein inference workflow set to maxLFQ, inference algorithm set to IDPicker, quantity level set to MS2, cross-run normalization set to false, and the protein grouping quantification set to median peptide and precursor quantity. Protein MS2 intensity values were assessed for quality using ProteiNorm (Graw *et al.*). The data was normalized using VSN (Huber et al) and analyzed using proteoDA to perform statistical analysis using Linear Models for Microarray Data (limma) with empirical Bayes (eBayes) smoothing to the standard errors (Thurman et al, Ritchie et al). Proteins with an FDR adjusted p-value < 0.05 and a fold change > 2 were considered significant.

**Acknowledgement:**
IDeA National Resource for Quantitative Proteomics and NIH/NIGMS grant R24GM137786.
